# CasDinG is an ATP-dependent 5’-3’ DNA helicase with accessory domains essential for type IV CRISPR immunity

**DOI:** 10.1101/2022.08.23.504870

**Authors:** Hannah Domgaard, Christian Cahoon, Matthew J. Armbrust, Olive Redman, Aaron Thomas, Ryan N. Jackson

## Abstract

CRISPR-associated DinG protein (CasDinG) is essential to type IV-A CRISPR function. However, the enzymatic activities of CasDinG are unknown. Here we demonstrate that CasDinG from *Pseudomonas aeruginosa strain 83* is an ATP- and metal-dependent 5’-3’ DNA helicase. The crystal structure of CasDinG reveals a helicase core of two RecA-like domains with three accessory domains (N-terminal, arch, and vestigial FeS). To examine the *in vivo* function of these CasDinG domains, we first identified the preferred PAM sequence (5’-GNAWN-3’ on the 5’-side of the target) with a plasmid library containing all combinations of the five nucleotides upstream of the target sequence. Plasmid clearance assays (using a 5’-GGAAA-3’ PAM) with CasDinG domain mutants demonstrated the vFeS and arch accessory domains are both essential for type IV immunity. These results provide a needed structural and biochemical framework for understanding the type IV-A CRISPR system.

## INTRODUCTION

CRISPR-Cas systems are prokaryotic immune systems that protect against mobile genetic elements, such as viruses and plasmids (1, 2). Of the six known CRISPR system types, type IV systems are the least understood and display a large diversity of protein composition (3, 4). Type IV-A systems rely on a multi-subunit complex consisting of a CRISPR RNA (crRNA) bound by several type IV-specific CRISPR-associated proteins (Csf1, Csf2, Csf3, Csf5/Cas6) and the CRISPR-associated DinG (CasDinG) protein to clear invasive plasmids (5–7) (Figure 1A). Additionally, mutations in the CasDinG Walker A or Walker B motifs impair immunity (6, 7), suggesting CasDinG mediated ATP binding and hydrolysis is essential for type IV-A function.

**Figure 1.**
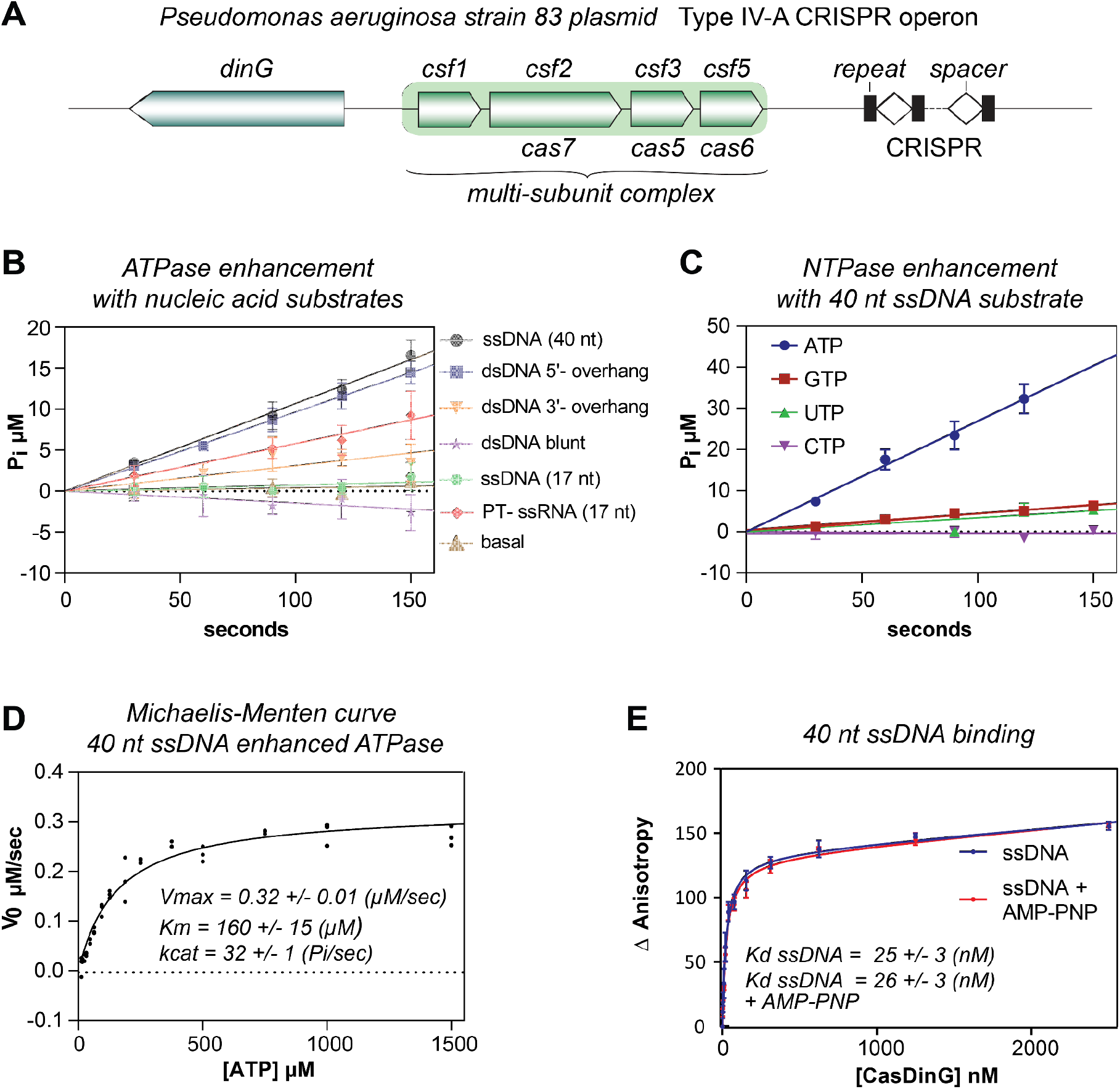
CasDinG is a nucleic acid-dependent ATPase. (**A**) Type IV-A CRISPR operon from *P. aeruginosa strain 83 plasmid*. (**B**) V_0_ plots depicting inorganic phosphate (P_i_) production via ATP hydrolysis with 10 nM CasDinG in the presence of 100 nM nucleic acid substrate and 600 µM ATP. (**C**) V_0_ plot depicting P_i_ phosphate production via NTP hydrolysis in the presence of 10 nM CasDinG and 600 µM NTP. (**D**) Michaelis-Menten curve depicting rates of P_i_ phosphate production at various ATP concentrations in the presence of 100 nM 40 nt ssDNA substrate with 10 nM CasDinG. (**E**) Fluorescence anisotropy curve of CasDinG binding to a 40 nt. 3’-FAM labeled ssDNA with and without AMP-PNP.

DinG (*damage-inducible gene G*) proteins are superfamily 2 helicases in the same family as eukaryotic XPD helicases involved in nucleotide excision repair (8–10). Deletion of the *dinG* gene in *E. coli* results in ultraviolet radiation sensitivity and mutations in XPD protein cause genetic diseases (11, 12). Notably, DinG proteins from different bacterial species display variability in domain organization and the functional role of ATP binding and hydrolysis. For example, in *E. coli*, DinG is a 5’-3’ DNA helicase that relies on ATP hydrolysis and an accessory domain containing an iron-sulfur (FeS) cluster to unwind substrates (12–14), while *S. aureus* DinG does not unwind DNA and lacks a FeS cluster domain (15). Instead, *S. aureus* DinG contains an N-terminal domain with 3’-5’ exonuclease activity that is regulated by ATP binding.

CasDinG proteins are distinct from non-Cas DinG sequences (16), including the *E. coli* and *S. aureus* DinG proteins (Supplementary Figure 1). For example, CasDinG contains an accessory domain in the same location as the *E. coli* FeS cluster domain, but the CasDinG domain lacks conserved cysteines for coordinating irons (16). Additionally, CasDinG has an N-terminal accessory domain, but it is smaller than the *S. aureus* N-terminal domain and it is not predicted to harbor exonuclease activity (Supplemental Figure 1). Thus, sequence comparisons to previously investigated bacterial DinG proteins could not fully reveal the biochemical activities of CasDinG.

To better understand the function of CasDinG in type IV-A CRISPR immunity, we performed helicase, ATPase, and nucleic acid binding assays with recombinant CasDinG protein encoded on a *Pseudomonas aeruginosa strain 83* extrachromosomal plasmid (NCBI ref: NZ_CP017294.1) (Figure 1A). We also solved the x-ray crystal structure of CasDinG, defined the type IV-A protospacer adjacent motif (PAM) consensus motif with a plasmid curing library, and performed cell-based assays with domain deletion mutants to explore the function of CasDinG accessory domains. Our results indicate CasDinG is an ATP and metal-dependent 5’-3’ DNA helicase containing an SF2 helicase core and three accessory domains. Two of these domains (vFeS and arch) are essential for type IV immunity. This work provides a structural and biochemical foundation for understanding the role of CasDinG in type IV-A CRISPR immunity.

## MATERIALS AND METHODS

### Construct generation & Recombinant Expression & Purification

The *dinG* gene from a *P. aeruginosa strain 83* extrachromosomal plasmid (NCBI ref: WP_088922490.1 (protein sequence), NZ_CP017294.1 (plasmid sequence)) was synthesized by TWIST bioscience and cloned into a pET StrepII TEV ligation independent cloning (LIC) vector (2R-T) vector (6). The annotated gene starts with a non-canonical (TTG) start codon, and the open reading frame continued upstream of the gene annotation. Thus, to ensure we were expressing the entire biologically relevant sequence we included the in-frame sequence upstream of the annotation starting with another non-canonical (TTG) start codon before reaching a stop codon. This approach added 48 DNA bases to the annotated gene and 16 additional amino acids (MKLAQGAFVDVIRIGA) to the N-terminus of the annotated amino acid sequence. Additionally, the original TTG start codon would likely encode for a leucine instead of methionine. The cloned vector was transformed into *Escherichia coli* BL21 HMS174(DE3) chemically competent cells (Novagen). A colony was picked and placed into an overnight outgrowth. In a 2.8 L flask, 1 L of Luria-Bertani (LB) medium supplemented with 1 mL of 1000 x metals mix (0.1 M FeCl_3_-6H_2_O, 1 M CaCl_2_, 1 M MnCl_2_-4H_2_O, 1 M ZnSO_4_-7H_2_O, 0.2M CoCl_2_-6H_2_O, 0.1 M CuCl_2_-2H_2_O, 0.2M NiCl_2_-6H_2_O, 0.1 M Na_2_MoO_4_-2H_2_O, 0.1 M Na_2_SeO_3_-5H_2_O,0 .1 M H_3_BO_3_) and 1 mL of 1000 x MgSO_4_ (1 M) was inoculated with 20 mL of overnight starter. Cells were grown to an optical density between 1.0 - 1.3 OD_600_ at 37 °C, then induced with a final concentration of 0.5 mM IPTG (isopropyl B-D-1-thiogalactopyranoside), while dropping the temperature to 20 °C. After 5 hours cells were harvested via high-speed centrifugation and stored at -80 °C.

Cells were homogenized on ice with lysis buffer (100 mM Tris Base pH 8.0, 150 mM NaCl, 1 mM TCEP). Protease inhibitors Aprotinin 1000 x (0.5mg/mL), Leupeptin 1000 x (0.5 mg/mL), Pepstatin A 1000 x (0.7 mg/mL), & PMSF (phenylmethylsulfonyl fluoride) 150 x (25 mg/mL) were added prior to cell lysis. Probe sonication for cell lysis was performed at settings of 4/60 (power/output). The lysate was clarified by high-speed centrifugation at 16,000 RPM for 30 minutes. All purification steps were performed at 4°C. The supernatant was loaded onto strep resin (Strep-Tactin®XT 4Flow®, IBA). The resin was washed with lysis buffer and then eluted with elution buffer (100 mM Tris Base, 150 mM NaCl, 50 mM biotin, 1 mM TCEP, pH 8.0). Fractions with CasDinG were pooled and desalted (HiPrep 26/10 Desalting, GE Healthcare) into low salt buffer (100 mM Tris Base pH 8.0, 10 mM NaCl, 1 mM TCEP). Elutions were then run over a heparin column (HiTrap Heparin HP, GE Healthcare), washed with 47.5 mM NaCl buffer and eluted with a high salt buffer (100 mM Tris Base pH 8.0, 500 mM NaCl, 1 mM TCEP). Samples were spin concentrated (Corning® Spin-X® UF 50 MWCO) before further purification with size exclusion (HiLoad 26/600 Superdex 200 pg., GE Healthcare) into the high salt buffer (100 mM Tris pH 8.0, 500 mM NaCl, 1 mM TCEP). Protein samples were concentrated and stored at 4°C. Protein was assessed for purification after each step via 12% SDS-PAGE. Notably, we observed that glycerol and/or freezing of the purified protein impaired CasDinG enzymatic activities.

Protein concentration was determined by UV-Vis spectroscopy (Thermofisher UV-vis Nanodrop), using the Beer-lambert law to correct absorbance values for extinction coefficient, wild-type 95910 M^-1^ cm^-1^ (assuming all cysteines are reduced), and molecular weight, 81131 daltons, as determined by Expasy Protparam(17).

### Clustal Omega Alignments and EMBOSS NEEDLE pair-wise alignments

*E. coli* (P27296) and *S. aureus* (Q2FGY5) DinG sequences were obtained through the Uniprot database server and aligned to Pa83 CasDinG using the Clustal Omega multiple sequence alignment tool (18). Pairwise sequence alignments were performed between *Pa83* CasDinG and *E. coli/S. aureus* sequences using an EMBOSS Needle alignment (19). Structural alignments between domains were either performed with the Secondary Structure Matching tool in Coot (20, 21), the DALI server (22) or the align command in the PyMOL Molecular Graphics System, Version 2.0 Schrödinger, LLC.

### Nucleic Acid Substrate Preparation

Nucleic acids were synthesized by Integrated DNA Technologies (IDT) (Supplementary Table 1). Nucleic acids were labeled with a fluorescein (FAM) label on the 5’ or 3’ end by IDT. To make duplexed nucleic acids, complementary oligonucleotides were mixed in an equimolar ratio in the presence of NEB buffer 2.1 and heated to 95°C. These oligonucleotides were slowly cooled to room temperature before being run on 12-15% NATIVE PAGE gels. Duplex bands were then gel extracted, ethanol precipitated, and reconstituted in water. For a list oligonucleotides used in assays, refer to Supplementary Table 1.

### Malachite Green ATPase Assays

Concentrations of P_i_ were determined with a Malachite Green Phosphate Assay kit (BioAssay Systems, Haward, CA, USA). Activated Malachite Green reagent was added to wells of a 384 well plate (Corning Assay Plate, 384 wells, Black with clear bottom, non-binding surface, Low flange, no lid, polystyrene, 3766). Before the reaction, assay components were pre-incubated at 37°C for 15 minutes. Reactions were started with the addition of ATP, and were run at 37°C. The reaction was quenched in the activated Malachite Green reagent at designated time points, between 30 - 150 seconds for reactions with activating nucleic acid and 1 - 4 minutes for basal reactions. The quenched reactions were developed for 30 minutes before sample measurement. The absorbance values of the samples were obtained using a Synergy H4 Hybrid Multi-Mode Microplate Reader measuring absorbance at 620 nm.

Comparison of initial velocities with various nucleic acid substrates utilized 10 nM CasDinG, 100 nM nucleic acid, and 600 µM ATP that was reconstituted in lab and spectroscopically verified, in the buffer (50 mM Tris pH 7.5, 0.1 mg/ml Recombinant Albumin, 1 mM MgCl_2_, 1 mM TCEP). Initial velocities of CasDinG NTP hydrolysis were determined in the presence of 10 nM CasDinG, 100 nM ssDNA (40 nt.) and 600 µM nucleotide triphosphate (NEB) in the buffer (50 mM Tris pH 7.5, 0.1 mg/ml Recombinant Albumin, 1 mM MgCl_2_, 0.4 µM TCEP). Michaelis-Menten curves were generated for CasDinG (20 nM) + single stranded phosphorothioated RNA (200 nM), and CasDinG (10 nM) + single stranded DNA (100 nM). Michaelis-Menten kinetic utilized the ATPase buffer (50 mM Tris pH 7.5, 0.1 mg/ml Recombinant Albumin, 1 mM MgCl_2_, 0.4 uM TCEP)

Initial velocities were performed with increasing concentrations of ATP (12-2000 µM), and a standard curve was generated using known concentrations of orthophosphate (0-100 µM). Initial velocities were calculated using linear regression. For a list of oligonucleotides used in these assays, see Supplementary Table 1. GraphPad Prism for Windows version 9.3.0 was used to fit the data using non-linear regression to the Michaelis-Menten equation (Equation 1). Where □ is the initial reaction velocity of the reaction, [S] is the ATP concentration, K_M_ is the Michaelis constant, and V_max_ is the maximum velocity of the enzyme.

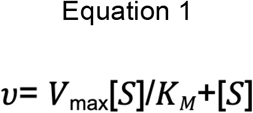

ATP concentrations were spectroscopically verified at 280 nm with crystal cuvettes and a spectrophotometer. Concentrations were confirmed using the Beer-Lambert law (Equation 2). Where A is the absorption, ε is the extinction coefficient in M^-1^cm^-1^, l is the path length in cm, and c is concentration in M. The molar extinction coefficient used for ATP was 15,400 M^-1^cm^-1^. ATP was frozen at -80°C, and aliquots were used and then discarded after a single thaw.

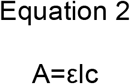

### Helicase assays

#### Substrate Comparisons

15 nM 5’ Fluorescein (FAM) labeled nucleic acid was incubated in the presence of 100 nM WT CasDinG and 1 mM ATP in the helicase buffer (25 mM Tris pH 7.5, 1 mM MgCl_2_, 1 mM TCEP, 0.1 mg/ml recombinant albumin) with 500 nM unlabeled displaced strand to protect against reannealing of displaced strands for approximately 20 minutes at 37°C before being quenched in 2x STOP Buffer (10 mM EDTA (Ethylenediaminetetraacetic acid), 1% SDS (Sodium Dodecyl Sulfate), 20% glycerol). Samples were run on 15% TBE NATIVE gels.

#### NTP Comparisons

15 nM 5’ FAM-labeled nucleic acid was incubated in the presence of 100 nM WT CasDinG and 1 mM ATP Analogue (ATP, ADP, ATPγS, and AMP-PNP) in the helicase buffer (25 mM Tris pH 7.5, 1 mM MgCl_2_, 1 mM TCEP, 0.1 mg/ml recombinant albumin) for approximately 20 minutes at 37°C before being quenched in 2x STOP Buffer with 500 nM of the unlabeled displaced strand. Samples were run on 15% TBE NATIVE gels

#### Metal Comparisons

15 nM 5’ FAM labeled nucleic acid was incubated in the presence of 100 nM WT CasDinG and 1 mM ATP and 1 mM divalent salt (MgCl_2_, MnCl_2_, ZnCl_2_, CaCl_2_, NiCl_2_, CoCl_2_, CuCl_2_) in the helicase buffer (25 mM Tris pH 7.5, 1 mM TCEP, 0.5 mg/ml BSA) for approximately 20 minutes before being quenched in 2x STOP Buffer with 500 nM unlabeled displaced strand. Samples were run on 15% TBE NATIVE gels

#### Time Courses

15 nM FAM-labeled nucleic acid substrate was exposed to 25 nM CasDinG over 10 minutes in the presence of 1 mM MgCl_2_ and 1 mM ATP in the helicase buffer (25 mM Tris pH 7.5, 0.1 mg/mL recombinant albumin, 1 mM TCEP) at 37°C. Samples were quenched in 2 x STOP Buffer at times between 0-10 minutes and run on 15% TBE native PAGE gels. The mutant analysis utilized the same nucleic acid substrate as WT CasDinG and an equivalent amount of protein. Samples were run on 15% TBE NATIVE gels.

#### Gel Analysis

All NATIVE PAGE gels for helicase assays were imaged using a BioRad Imaging system and analyzed using BioRad ImageLab software. Percent unwound were quantified using ImageLab software and the reported data is the average of three experiments, with error bars representing the standard deviation from the mean. Graphs were made in GraphPad Prism for Windows version 9.3.0.

#### Nucleic acid-binding assays

Nucleic acid-binding activities of strep-tagged CasDinG were monitored using a fluorescence polarization-based assay. Anisotropy data were collected using a BioTek Synergy H4 Hybrid Multi-Mode Microplate Reader equipped with polarizers and bandpass filters. The polarizers and bandpass filters provided 485 ± 20 nm excitation and detection of fluorescence emission at 528 ± 20 nm. Each reaction (80 µL) contained a limiting concentration (10 nM) of 3’ FAM-labeled ssDNA (40 nt) and 5’ FAM-labeled ssDNA (17 nt). CasDinG and 3’-end FAM labeled nucleic acid were assayed at room temperature with increasing concentrations of CasDinG (0 – 2.5 µM) in a binding buffer (100 mM Tris pH 8.0, 1 mM TCEP, and 5 mM MgCl_2_) with or without 1 mM AMP-PNP (Sigma Aldrich). Change in anisotropy relative to FAM-nucleic acid was plotted as a function of CasDinG concentration. The apparent dissociation constant (K_d_) for the nucleic acid substrate was determined by fitting the raw data to a single site saturation binding model in GraphPad Prism for Windows version 9.3.0.

#### Crystallization and Structure determination

Strep-tagged CasDinG protein was concentrated to 5 mg/ml and crystallized using 225 mM Imidazole pH 8.0, 3.5% PEG 8000, and 30% sucrose with hanging-drop vapor diffusion at room temperature. The crystal used for structure determination was comprised of 1 uL (5mg/ml) protein solution to 2.6 uL mother liquor and 0.4 uL 30% sucrose. The crystal was then soaked in a cryoprotecting solution composed of 30% Ethylene glycol and mother liquor, then mounted on a loop, and cooled to 100 K. Diffraction data were collected at the SSRL beamline 9-2. The data were indexed, integrated, and scaled using HKL3000 to 2.95 Å resolution with the space group P65 (23). Phases were determined by molecular replacement in Phaser (24, 25) with an N-terminally truncated Alphafold structure prediction with Colab (26), prepared with the Process Predicted Model tool in Phenix to adjust the model B-factors (27). After obtaining an initial solution with Phaser, RESOLVE was used to obtain a density-modified map (28). Because the space unit cell consisted of ∼70% solvent, density modification significantly improved the electron density maps. Model building was performed in Coot (21), structures were refined using PHENIX, and validation was performed using Molprobity within PHENIX and the PDB deposition servers (27).

#### Preparation of electrocompetent cells for PAM assay

50ul of chemically competent HMS174(DE3) cells were first doubly transformed with roughly 20 ng each of plasmids #1290 and #1284 (targeting), and another vial of HMS174(DE3) cells was also transformed with #1291 and #1284 (control). Cells were transformed by heat shock at 42°C followed by a cold shock on ice for 1-3 minutes. 450 µL of SOC was then added to each transformation, followed by a 1hr recovery period in a shaking incubator at 37°C. 2 dilutions of each tube were then plated on double antibiotic LB agar plates (chloramphenicol 25 µg/mL, streptomycin 50 µg/mL). Plates were then allowed to grow overnight at 37°C, and the next day 3 well isolated colonies were picked from both a targeting plate and a control plate and each was used to inoculate a 5 mL overnight growth of LB with selection by chloramphenicol and streptomycin. Overnight growths were carried out in a shaking incubator at 37°C.

The following day, 2 mL of each overnight growth was used to inoculate 48 mL of fresh LB with selection antibiotics, and the culture was then allowed to grow until reaching an OD of approximately 0.25. At this point IPTG was added to a concentration of 0.1 mM to induce expression of the CRISPR-Cas system, and the cells were allowed to grow for another 50 minutes. The cultures were then transferred to 50 mL conical tubes and spun down at 4000 x gat 4°C for 15 minutes to pellet them. Pellets were then resuspended in 50ml of ice-cold sterile MilliQ filtered water followed by another round of centrifugation at 4000 x g at 4°C for 15 minutes. Pellets were then resuspended in 2 mL of ice-cold sterile 10% glycerol in a new 15 mL conical tube and spun down again at 4000 x g at 4°C for 10 minutes. Cells were resuspended one more time in 460 µL of ice-cold 10% glycerol and used immediately for electroporation.

#### *In Vivo* PAM assay

A lyophilized PAM library in the form of a pET backbone containing a CA01 target sequence flanked on the 5’ end by a library of all potential base combinations of the five positions immediately upstream of the crRNA target was provided by the Chase Biesel laboratory at the Helmholtz Center for Infection Research, and resuspended in nuclease-free water to a concentration of roughly 500 ng/µL. 1.5 µL of the PAM library was then added to 100 µL of a vial of fresh electrocompetent cells containing either a functional (targeting) or non-functional (control) immune system. This was done in biological triplicate with 3 targeting and 3 control transformations. The cells were then transferred to a pre-chilled 2 mm electroporation cuvette. Recovery media consisting of SOB with 0.1 mM IPTG was prepared for each transformation. Electroporation was carried out for each cuvette with settings of 2500V, 25 µF, and 400Ω, with a typical time constant of 4-5 ms. Immediately after applying the voltage 1 mL of recovery media was added, and the cells were transferred to 1.5 mL microcentrifuge tubes to recover in the incubator at 37°C with shaking.

After recovery, 1ml of each transformation was used to inoculate 50 ml of LB with triple antibiotic selection (chloramphenicol 25 µg/mL, streptomycin and kanamycin 50 µg/mL). Liquid cultures were then allowed to grow overnight for 20 hours at 37°C with shaking, and then harvested by centrifugation at 4000 x g at 4°C for 15 minutes. Plasmid DNA was then extracted from each pellet with an Omega Bio Tek E.Z.N.A.® Plasmid DNA Midi kit.

#### Next-Generation Sequencing

Primers flanking the insertion site in the plasmid were used to amplify the PAM sequences from the PAM library from each of the 3 target and 3 control samples. To these primers were added the Illumina Truseq sequencing primer sequences. This reaction was performed for 22 cycles using Q5 (New England Biolabs). These Illumina sequences were then used as the template for a second of PCR to add indexes and the p5 and p7 Illumina adapter sequence to make finished sequencing libraries. This PCR was performed for 10 cycles using Q5 These libraries were then sequenced on an Illumina MiSeq sequencer using the 300 cycle v2 chemistry. After demultiplexing approximately 500,000-600,000 reads were obtained for each sample.

#### PAM sequence analysis

A short script using a regular expression was used to parse the 5 nucleotides preceding a correct target sequence from each read, and the number of occurrences of each potential NNNNN PAM was counted and exported as a CSV file. Counts were summed for all targeting and control runs, and a depletion score was calculated for each PAM sequence which relates the proportion of all counts of a PAM in the targeting runs to the control runs.

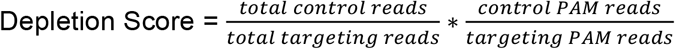

All PAM sequences along with their depletion scores were then input into a Krona plot generator (29) which was in turn used to generate a PAM wheel.

#### Preparation of chemically competent Type IV-A CRISPR-Cas cells

HMS174(DE3) *E. coli* cells (Novagen), were transformed with 50-100 ng of both plasmid #1284(pCDF_Pa_csf1_csf2_cas6) and plasmid #1290(pACYC-PaCR83-csf3-DinG) containing WT CasDinG, or plasmids containing the mutant, #2368 (PACYC-PA83CR-csf3-DinG-ArchKO), #2369 (PACYC-PA83CR-csf3-DinG-FeS-KO), or #2370 (PACYC-PA83CR-csf3-DinG-DEAH_AAAH-KO). Mutants were generated using the Q5 Mutagenesis kit (NEB Biolabs).

Colonies from the overnight plates were used to inoculate 25 mL of the prepared LB broth in a 50 mL Falcon tube. These cells were then incubated at 37°C until an OD_600_ between 0.2-0.3 when they were induced with 0.1 mM IPTG (100 µM IPTG). Cells were allowed to grow for an additional 45 minutes at 37°C before being cold-shocked on ice for 20 minutes. Cells were then spun down at 2700 x G for 15 minutes at 4°C. The supernatant was decanted, and the cells were resuspended in 12 mL RF1 Buffer (100 mM RbCl, 50 mM MnCl_2_4H_2_O, 30 mM Potassium acetate, 15% m/v glycerol). Cells were then allowed to rest on ice for 15 minutes before being spun down at 870 x G for 15 minutes. The supernatant was decanted and cells were resuspended in 1 mL of RF2 Buffer (10 mM MOPS, 10 mM RbCl, 75 mM CaCl_2_2H_2_O 15% m/v glycerol). Cell solution was incubated on ice for 15 minutes prior to aliquoting cells in 100 uL volumes and flash freezing at -80 °C.

#### *In Vivo* Plasmid Competition Assay

Using type IV-A CRISPR-Cas containing chemically competent cells, 10 ng of target or non-target plasmid, #2380 (pET27b_CA01_GGAAA) and #1095 (pET27b(+)-non-target_TTTC) respectively, was added to a 100 uL cell aliquot. Cells were then heat shocked at 42°C for 30-40 seconds, followed by cold shock on ice for 1-3 minutes. 400 µL of LB containing chloramphenicol 25 µg/mL, streptomycin 50 µg/mL, and 0.1 mM IPTG were then added to the cold shocked cells. Cells were then incubated at 37 °C for 45-50 minutes in a shaking incubator, 200 RPM, followed by plating of cells onto a triple antibiotic LB agar selection plate (Chloramphenicol 25 µg/mL, streptomycin and kanamycin 50 µg/mL, and IPTG 0.1 mM IPTG). 400 uL of cells were plated onto one plate followed by 100 µL on another for a □ and □ dilution respectively. The cells were then spread around by shaking with 3 sterile glass beads per plate and subsequently allowed to dry before being placed in a 37 °C incubator for 24 hours. Colonies were then counted manually.

## RESULTS

### CasDinG is a nucleic acid-dependent ATPase

The Walker A and B motifs of CasDinG were recently shown to be essential for type IV immunity (6, 7). Although the canonical function of these motifs is ATP binding and hydrolysis, it was unclear if CasDinG uses ATP to unwind DNA duplexes, like *E. coli* DinG (12) or to regulate accessory activities, like *S. aureus* DinG (15). Thus, to determine the role of ATP in CasDinG function we expressed and purified recombinant CasDinG from *P. aeruginosa strain 83* extrachromosomal plasmid (Supplementary Figure 2), and used a malachite green assay to detect the release of inorganic phosphate from ATP (Figure 1B). Because ATPase activities are often allosterically enhanced by the presence of nucleic acid in helicases (30, 31), we examined ATP hydrolysis in the presence of various nucleic acid substrates (Figure 1B). We observed that a 40 nt. ssDNA, 17 nt. phosphorothioate RNA (PT-ssRNA), and 17 nt. DNA duplex with a 17 nt. 5’-overhang all strongly enhanced ATPase activity. A 17 nt. DNA duplex with a 17 nt. 3’-overhang substrate moderately enhanced ATPase activity, while the 17 nt. blunt duplex and 17 nt. ssDNA did not enhance ATPase activity above background. To evaluate whether CasDinG can hydrolyze other ribonucleotide triphosphates we measured the release of inorganic phosphate from ATP, GTP, CTP and UTP (Figure 1C). CasDinG hydrolyzed GTP and UTP, but at a rate fourfold lower than ATP hydrolysis, and no CTP hydrolysis was observed, indicating CasDinG preferentially hydrolyzes ATP.

**Figure 2.**
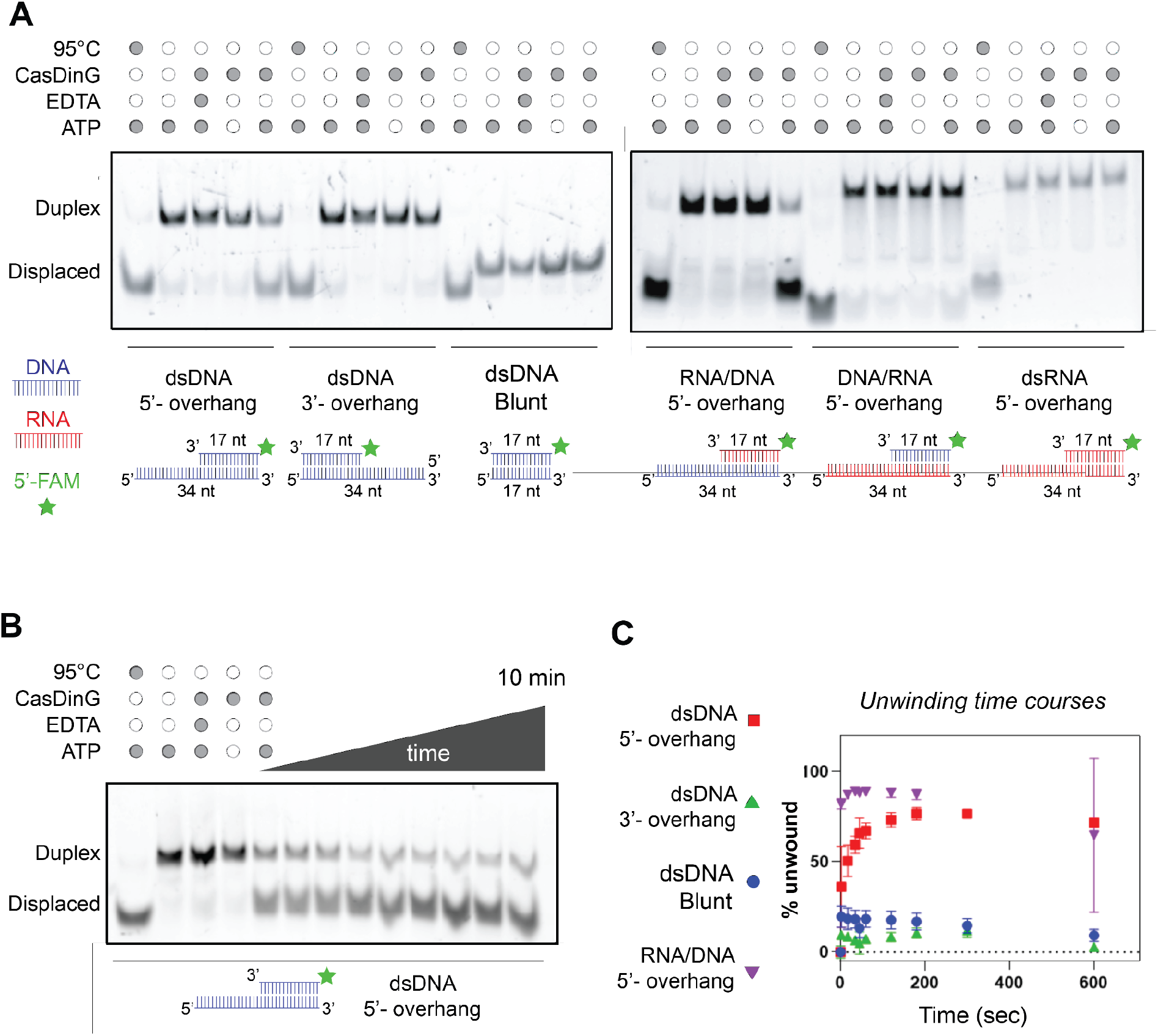
CasDinG is a 5’-3’ ATP and metal dependent helicase. (**A**) Native PAGE helicase assay showing displacement of a FAM-labeled ssDNA or ssRNA from a duplexes with a 5’-DNA overhang. Minimal to no displacement is visualized for 3’ overhangs, blunt ended duplexes, or substrates in which the 5’ overhang is RNA. All reactions were quenched after 20 minutes (**B**) Native PAGE helicase assay showing a time course unwinding of 5’-FAM labeled dsDNA 5’-overhang. The reaction took place over the course of 10 minutes using 10 nM CasDinG and 15 nM FAM labeled substrate. (**C**) A graph plotting the % of displaced FAM-labeled nucleic acid over time. Only the dsDNA 5’overhang and RNA/DNA hybrid 5’ DNA overhang unwound more than 25%.

To further evaluate how nucleic acid enhances ATPase activity, we collected ATP hydrolysis velocities with 40 nt ssDNA and 17 nt ssPT-RNA at different concentrations of ATP and fit the values to the Michaelis Menton equation (Figure 1D and Supplementary Figure 3). We observed that apo CasDinG only weakly hydrolyzed ATP with an observable velocity of < 0.2 µM sec^-1^ at the highest concentration of ATP used (2 mM) with 50 times more enzyme than assays with nucleic acid (500 nM). This activity was enhanced at least 100-fold and 75-fold by a 40 nt. long ssDNA (k_cat_ of 32 +/- 1 sec^-1^) or 17 nt. long ssPT-RNA (k_cat_ of 23 +/- 0.7 sec^-1^), respectively (Supplementary Figure 3), and similar to the k_cat_ of 24 sec^-1^ reported for *E. Coli* DinG in the presence of ssDNA (12).

**Figure 3.**
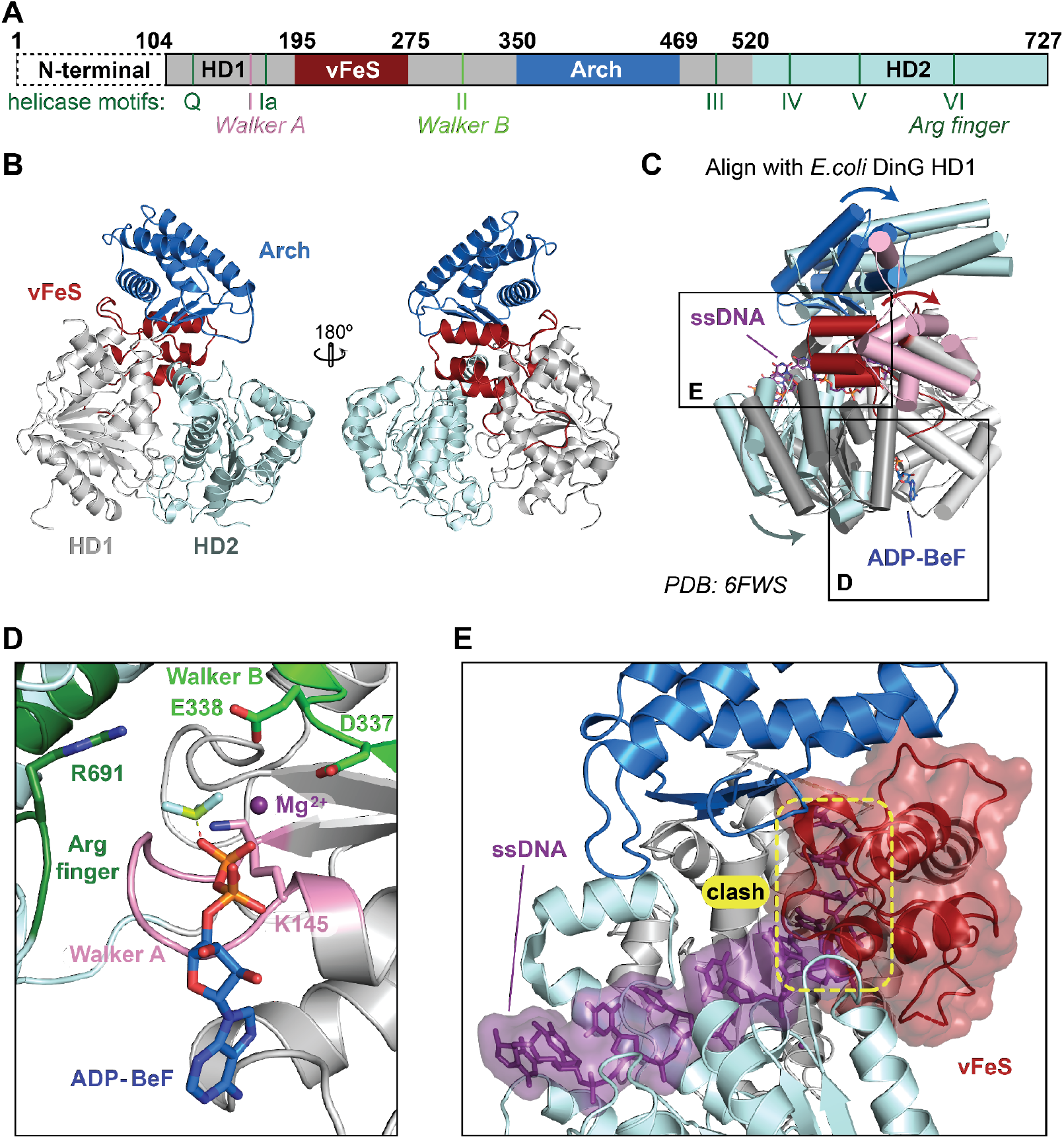
X-ray crystal structure of CasDinG. (**A**) Diagram of the CasDinG protein sequence depicting the HD1 (silver) and HD2 domains (pale cyan) with accesory domains vFeS (red) and Arch (blue). Position of SF2 helicase motifs are indicated under the diagram. The N-terminal domain of CasDinG is indicated but was unresolved in the solved structure. (**B**) Model of CasDinG highlighting the Arch and vFeS accesory domains. (**C**) Alignment of CasDinG model with the *E. Coli* DinG model bound to ssDNA and ADP-BeF3 Mg^2+^ complex (PDB 6FWS). Positional differences between the CasDinG and E. coli DinG are indicated with arrows (**D**) Zoomed in view of the ATP binding pocket with the aligned ADP-BeF3 Mg^2+^ complex (6FWS) to the CasDiG model. The Walker A (pink) and Walker B (lime) motifs are in position ot coordinate the ADP-BeF3 Mg^2+^. (**E**) View of the nucleic acid binding pocket showing the aligned ssDNA (6FWS) within the RecA-like fold of the HD1 and HD2 domains. Highlighted in yellow is a clash between the aligned ssDNA and the CasDinG vFeS domain, indicating the vFeS domain is in a closed position that would have to open to accommodate bound ssDNA.

Just as the presence of nucleic acid enhances ATPase activity, helicase affinities for nucleic acid substrates are often enhanced by the presence of ATP (32–34). To investigate if ATP enhances the ability of CasDinG to bind nucleic acid substrates we determined the dissociation constant (K_d_) of binding of a 40 nt. long ssDNA to CasDinG with and without the non-hydrolyzable analog AMP-PNP (Figure 1E). CasDinG bound to the ssDNA with a K_d_ of 25 +/- 3.1 nM without AMP-PNP, and with a K_d_ of 26 +/- 2.9 nM with AMP-PNP, demonstrating the presence of a nucleotide triphosphate does not enhance nucleic acid binding. Consistent with this observation, when the length of the DNA was shortened to 17 nt, the affinity of CasDinG to nucleic acid only slightly increased in the presence of AMP-PNP and ADP from a K_d_ of 81 +/- 5.4 nM to 47 +/- 4.9 nM and 37 nM +/- 3.2 respectively (Supplementary Figure 4). Collectively, these data indicate CasDinG is an ATP hydrolase that is activated by ssDNA or ssRNA. However, CasDinG affinity for nucleic acids is not dependent on nucleotide binding.

**Figure 4.**
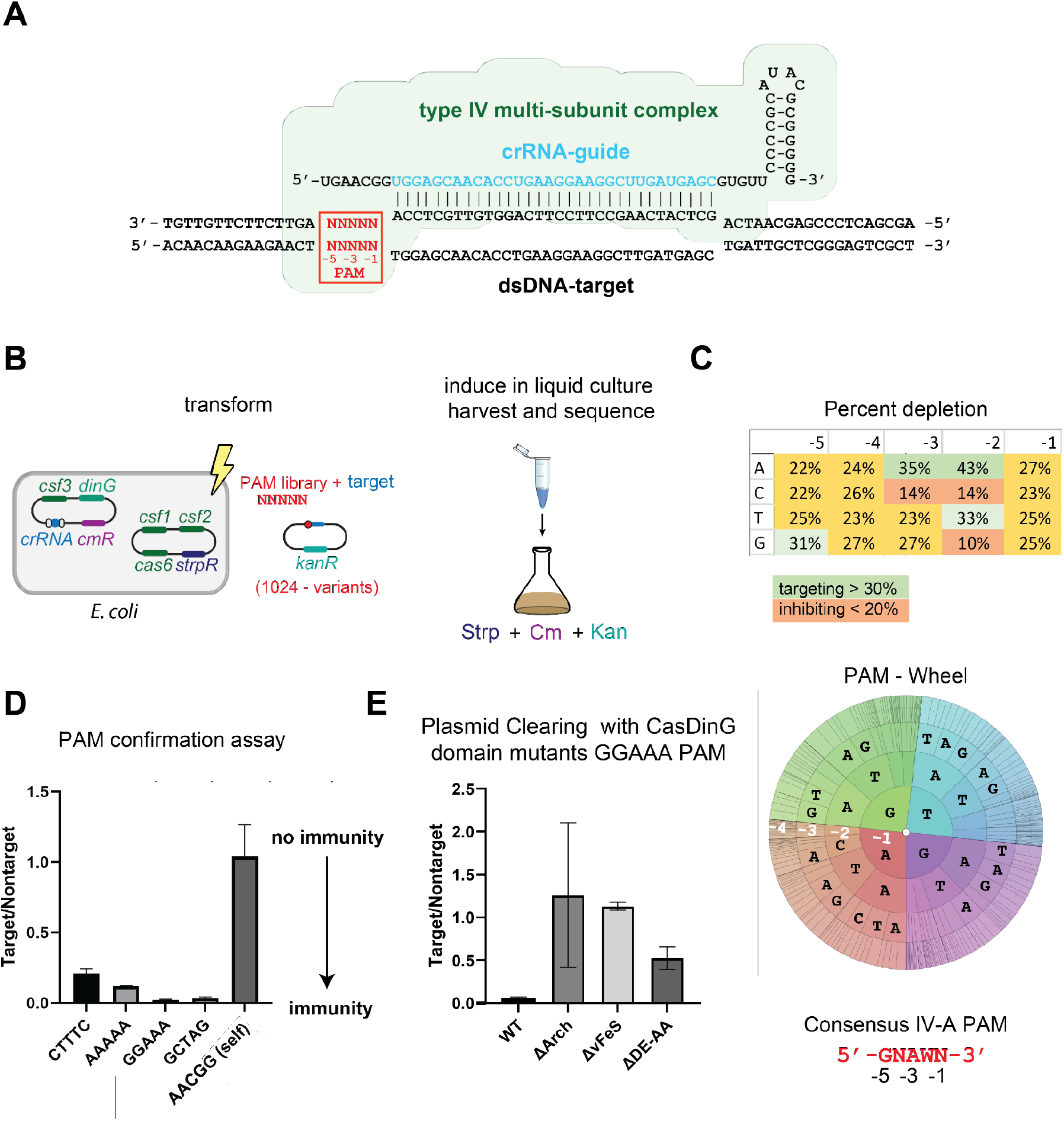
The type IV-A CRISPR system prefers a GNAWN PAM and requires CasDinG accesory domains. (**A**) Schematic depicting the type IV-A Csf complex binding to a dsDNA target. The location of the PAM is indicated in red. (**B**) Diagram of how the PAM library was performed. E. coli cells expressing the type IV-A system were transformed with a library of target plasmids adjacent to all possible combinations of PAM sequences. The transformation was grown up under antibiotic selection, cells were harvested, DNA was isolated and deep sequenced. (**C**) Table (top) and PAM wheel (bottom) showing percent depletion of nucleotides in the PAM assay, highlinghting a consensus PAM 5’-GNAWN -3’ sequence. (**D**) Ratio of target/nontarget colonies using four different targeting PAM sequences compared to self. (**E**) Plasmid curing assay with vFeS and Arch domain mutants, demonstrating thes domains are essential for type IV-A CRISPR immunity.

### CasDinG is an ATP- and metal-dependent 5’-3’ dsDNA helicase

Although we determined that CasDinG binds nucleic acid substrates and hydrolyzes ATP, it remained unclear if CasDinG uses these activities to unwind nucleic acid duplexes. To determine if CasDinG possesses helicase activity we combined CasDinG with various DNA duplexes. One strand of each duplex was 5’-end-labeled with fluorescein (FAM) to visualize strand displacement with an electromobility shift assay (EMSA) on a polyacrylamide gel. We observed that CasDinG displaced the FAM-labeled strand from 5’-overhang DNA duplexes, but not duplexes with a 3’-overhang or no overhang (blunt), consistent with a 5’-3’ unwinding polarity (Figure 2A). Additionally, CasDinG displaced RNA from DNA-RNA hybrid duplexes with 5’-DNA overhangs, but could not displace DNA or RNA from duplexes with 5’-RNA overhangs (Figure 2B). This observation suggests the helicase activity of CasDinG is DNA specific.

The helicase activities of CasDinG were only observed in the presence of ATP and were impaired by EDTA, indicating unwinding is coupled to ATP hydrolysis and requires a divalent metal. To confirm ATP hydrolysis is coupled to unwinding we examined helicase activity in the presence of ADP and non-hydrolyzable analogs ATP□S and ATP-PNP. Only in the presence of ATP was strong duplex unwinding observed (Supplementary Figure 5A-B). To determine which divalent metals activate CasDinG helicase activity, we examined helicase activity with a variety of divalent metals. We observed that Mg^2+^, Mn^2+^, Ca^2+^, Ni^2+^, and Co^2+^, allow for helicase activity, whereas larger cations Zn^2+^ and Cu^2+^ did not (Supplementary Figure 5C). Thus, CasDinG is an ATP and divalent metal ion-dependent 5’-3’ helicase capable of displacing RNA or DNA from a DNA substrate.

**Figure 5.**
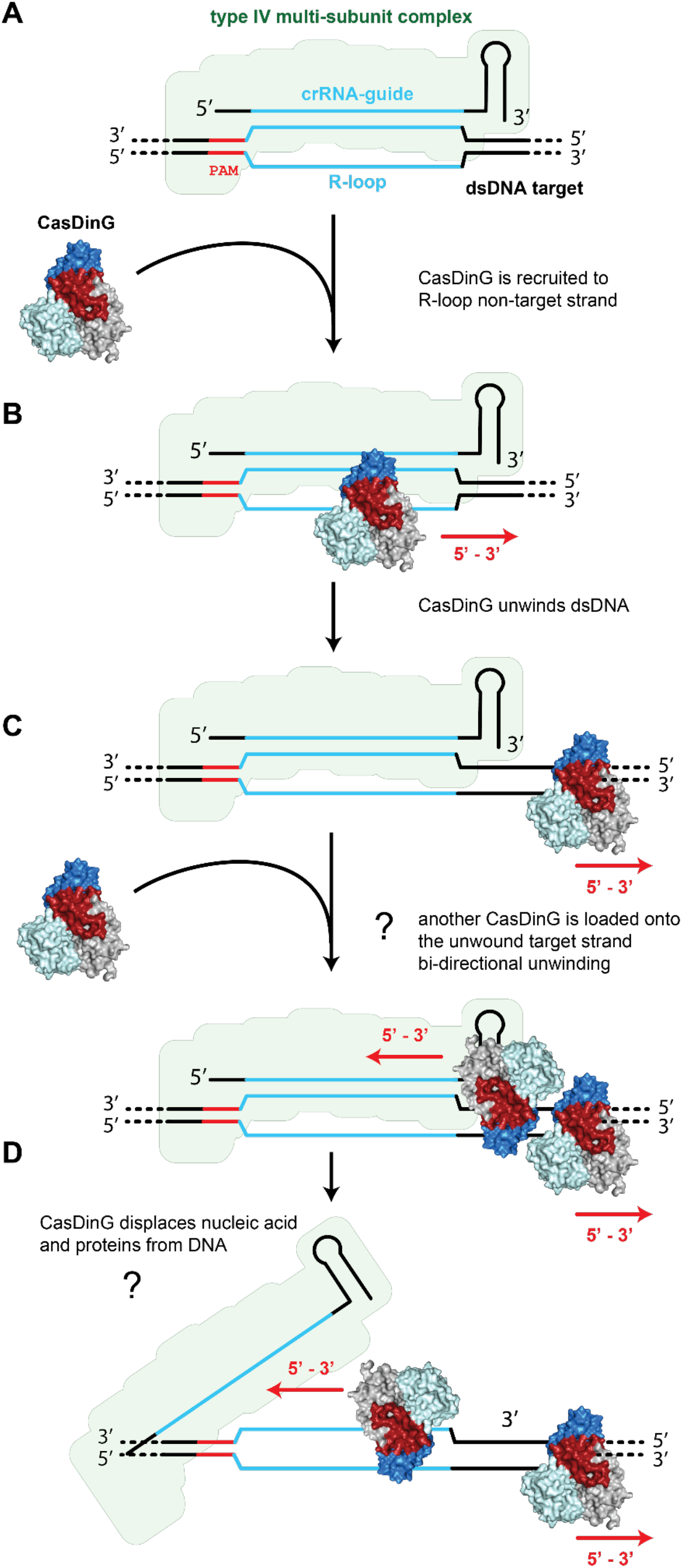
Model of the role of CasDinG in type IV-A CRISPR immunity. (**A**) The type IV-A multi-subunit Csf crRNA-guided complex binds complementary dsDNA forming an R-loop to which CasDinG is recruited (**B**) CasDinG unwinds the dsDNA in a 5’-3’ manner. (**C**) Another CasDinG may be loaded on the displaced target strand. (**D**) CasDinG unwinding perhaps displaces proteins, and primers involved in replication and/or transcription machinery.

### CasDinG structure consists of a helicase core with three accessory domains

CasDinG shares less than 22% sequence identity and 31% sequence similarity with both *S. aureus* and *E. coli* DinG (Supplemental figure 1). The highest similarity regions are the SF2 helicase motifs (Q, I, II, III, IV, V, VI) within the two RecA-like domains of the helicase core, while the least similar regions are within the predicted accessory domains (Supplementary Figure 1). We hypothesized that sequence differences in the accessory domains might influence the function of CasDinG in type IV-A immunity. To better understand the function of the accessory domains we solved the crystal structure of CasDinG from *Pseudomonas aeruginosa strain 83* at 2.95 Å resolution with an Rwork/Rfree of () in space group P 6_5_ (Figure 3 and Table 1) by molecular replacement (24, 25), and an alphafold model of N-terminally truncated CasDinG. Notably, the domains of the alphafold prediction aligned to our final model with RMSDs < 1 Å (Supplementary Figure 6). The only major differences between the alphafold model and our final structure were interdomain positions.

**Table 1.** Data collection and refinement statistics.

| Dataset | Native |
| --- | --- |
| **Data collection** |  |
| Beamline | SSRL 9-2 |
| Space group | $P6_5$ |
| Cell dimensions |  |
| *a*, *b*, *c* (Å) | 123.3, 123.3, 136.9 |
| $\alpha$, $\beta$, $\gamma$ (deg) | 90.0, 90.0, 120 |
| Wavelength (Å) | 0.88684 |
| Resolution<sup>†</sup> (Å) | 50-2.95 (3.06-2.95)* |
| *R*<sub>merge</sub>(%) | 18.8 (135.6) |
| CC<sub>1/2</sub> | 0.994 (0.766) |
| *I* / σ*I* | 20.0 (2.0) |
| Observations | 49009 (4849) |
| Unique reflections | 24916 (2453) |
| Multiplicity | 20.5 (16) |
| Completeness (%) | 100.0 (99.8) |
| Resolution (Å) | 50-2.95 (3.06-2.95) |
| No. reflections | 24916 |
| **Refinement** |  |
| *R*<sub>work</sub>/*R*<sub>free</sub> (%) | 18.5/21.5 |
| No. atoms |  |
| Protein | 4578 |
| Water | 0 |
| Ligands | 0 |
| B-factors |  |
| mean | 76.2 |
| R.m.s. deviations |  |
| Bond lengths (Å) | 0.002 |
| Bond angles (deg) | 0.52 |
| Ramachandran |  |
| Favored (%) | 96.33 |
| Allowed (%) | 3.77 |
| Outliers (%) | 0 |
| Clashscore | 3.4 |

\*Values in parentheses are for highest-resolution shell.
<sup>†</sup>Resolution limit used the criterion of *I*/ $\sigma$ *I* > 2.0

The CasDinG structure reveals a SF2 helicase core of two RecA-like helicase domains (HD1 and HD2) with three accessory domains; an N-terminal domain of 115 amino acids not observed in the electron density, and two inserts within HD1 (vFe/S and arch domain) (Figure 3A). Similar to other SF2 helicases, the conserved helicase motifs decorate the cleft between HD1 and HD2 (33) (Supplementary Figure 7). Alignment of the RecA helicase domains of CasDinG with *E. coli* DinG bound to ADP-BeF and ssDNA (PDB: 6FWS) shows that these enzymes are highly similar within the helicase core, and suggests CasDinG uses a similar mechanism as *E. coli* DinG to bind and hydrolyze ATP (Figure 3C and 3D). The alignment also suggests that DNA binds and translocates along the RecA lobes opposite the ATP binding site. However, in our structure the vFeS domain appears to be in a closed conformation, sterically blocking the path of the traversing DNA (Figure 3E), indicating rearrangement of the vFeS domain must occur to allow for nucleic acid binding and duplex unwinding.

Although the two CasDinG domains inserted within HD1 have low sequence similarity to *E. coli* DinG they share some remarkable tertiary topology (Supplementary Figure 8). The first insert (residues 195-275) consists of a four alpha-helix bundle, that aligns with the *E. coli* DinG FeS cluster domain using the Coot Secondary Structure Matching tool with an RMSD of 2.3 Å (20, 35). Despite this conserved arrangement, CasDinG lacks three of the four cysteines observed in *E. coli* DinG that coordinate the FeS cluster, and no FeS cluster is observed in the CasDinG electron density (Supplemental Figure 8A). Presumably, the FeS cluster coordination in *E. coli* DinG stabilizes the tertiary fold of the domain, holding α2 into proximity to the loop directly downstream of α4 (supplemental figure 8A). In CasDinG, the loss of stability provided by FeS cluster coordination is compensated for by a salt bridge formed between residues R204 and D269. These residues appear to be fairly conserved in CasDinG sequences but not in chromosomal DinG (16). Thus, to distinguish the CasDinG domain from DinG-like helicases with accessory domains that coordinate FeS clusters, we name this CasDinG accessory domain a vestigial FeS domain or vFeS. Additional differences between CasDinG and *E. coli* DinG in this domain are CasDinG lacks a loop-helix-loop connecting helices α2 and α3 of the *E. coli* DinG FeS domain, but CasDinG contains an extended loop helix connecting alpha helices α3 and α4 (Supplementary Figure 8). This extended loop sits within the RecA domain cleft suggesting it may help regulate ATPase or unwinding activity.

The second CasDinG accessory domain (residues 352-467) consists of a four-helix-bundle and a three-stranded anti-parallel beta-sheet, sharing structural topology with the arch domain of *E. coli* DinG and XPD helicases (35–37). Indeed, the DALI server reports an RMSD of 3.3 Å between aligned CasDinG and *E. coli* DinG arch domains (22). However, the arch domain of *E. coli* DinG is 61 amino acids larger and contains an extra beta-loop connecting helices α2 and α3. The arch domain of XPD makes important contacts with other proteins in the human transcription factor IIH complex (37, 38). Thus, it is possible that the sequence and size differences of the arch domains of CasDinG and *E. coli* DinG could be associated with differences in protein-protein interactions that these domains may mediate.

### The type IV-A system recognizes a 5’-GNAWN-3’ PAM

To better understand the function of the CasDinG accessory domains we desired to examine domain deletion mutants with the cell-based assay we previously developed to demonstrate type IV-A plasmid clearance (6). However, we were concerned that our assay was not optimal because recent literature suggested type IV-A systems prefer an 5’-AAG-3’ protospacer adjacent motif (PAM) located on the 5’-side of the target sequence (4, 7), instead of the 5’-CTTTC-3’ PAM our assay utilized. PAMs are small nucleotide motifs observed next to the nucleic acid targets of crRNA-guided surveillance complex that distinguish “non-self” from “self” targets (39–41).

To determine what PAM sequences optimally activate the type IV-A system, we performed a plasmid curing assay in duplicate with a library containing all 1024 combinations of 5 nucleotides adjacent to the 5’-side of the target sequence, similar to previous studies identifying PAM preferences (42) (Figure 4). Transfected cells were grown in liquid culture under immune system-inducing conditions, harvested, and deep sequenced. The lack or depletion of a PAM sequence, when compared to a no-immune system control, indicated an activating PAM sequence. Depletion scores for each PAM base position, and a PAM wheel (Figure 4C) were generated to reveal PAM nucleotide preferences. The data revealed a depletion preference for guanine at position -5, adenosine at position -3, and adenosine or thymine at position -2. There appeared to be no preference for nucleotides at positions -1 and -4, defining a preferred consensus PAM of 5’-GNAWN-3’, where W indicates A or T. Notably, there also appears to be an anti-targeting effect when guanine or cytosine is located at position -2, and when cytosine is located at position -3. Interestingly, the repeat region adjacent to the 5’-side of the spacer (self-sequence) contains a cystosine at this -3 position and a guanine at this -2 position, suggesting the PAM recognition mechanism has evolved to avoid self targets while gaining a mechanism to interrogate non-self targets.

To confirm that the 5’-GNAWN-3’ PAM is preferred we performed plasmid clearance assays with single target plasmids containing specific PAMs (Figure 4D). We first tested the 5’-CTTTC-3’ PAM used in our previous work. Although, this PAM does not contain an adenosine at the -3 position or a guanine at the -5 position we still observed depletion of the target strand compared to the non-target. However, consistent with our PAM library screen, PAMs that conformed better to the 5’-GNAWN-3’ consensus sequence, with either a -3 adenosine, a -5 guanine, or both (e.g. 5’-GGAAA-3’) showed stronger target clearance. Collectively, these data suggest that the reason the 5’-CTTTC-3’ PAM worked previously is because is did not contain anti-targeting bases G-C or C in the -2 or -3, position. However, other PAMs that conform to the 5’-GNAWN-3’ sequence are more preferred.

### CasDinG vFeS and arch domains are essential for type IV-A immunity

After identifying the optimal PAM for the type IV-A system, we used our cell-based assay to investigate the role of the CasDinG accessory domains in type IV immunity. We made deletions in the plasmid encoding CasDinG that deleted the vFeS or arch domain (Supplementary Figure 9) and used a target plasmid with a 5’-GGAAA-3’ PAM. We observed that deletion of either the vFeS or arch domain disrupted the immune system similar to the Walker B mutant, indicating these domains play an essential role in type IV-A immunity.

## DISCUSSION

CasDinG is an essential component of the type IV-A CRISPR Cas system, which clears bacteria of invasive targets and silences gene expression (6, 7). The type IV-A multi-subunit complex or Csf complex is presumed to bind to DNA targets, similar to the type I Cascade complex, and then recruit the CasDinG helicase onto the DNA target (5, 7)(Figure 5A). This presumption is supported by (i) recent work in *Pseudomonas oleovorans* demonstrating type IV-A systems silence LacZ expression when targeting the gene on either the coding or non-coding strand (7), and (ii) this work demonstrating CasDinG is a DNA, but not an RNA, helicase.

Previous bioinformatic and cell-based assays suggested the Csf complex prefers to target sequences downstream of a 5’-AAG-3’ PAM (4, 7). Here we used a target library to further define the PAM preference for DNA targeting as a 5’-GNAWN-3’ PAM with a strong resistance to targeting self sequences containing a cytosine in the -3 position or guanine or cytosine in the -2 position of the PAM. Notably, the previously reported 5’-AAG-3’ PAM conforms to this broader description, but some individually tested PAMs in the *P. oleovorans* system that fit our proposed consensus sequence were shown to be non-targeting (7). This observation could be due to unfavored bases in the -5 position that were not considered, or species-specific differences in the type IV-A PAM recognition mechanism. It should also be noted that our assay only looked as far as the -5 position. It remains possible that positions further downstream contribute to the self-vs-non-self PAM recognition mechanism.

Different class 1 multi-subunit CRISPR systems use distinct PAM recognition mechanisms to distinguish self from non-self targets (40). The type III systems, which target ssRNA, predominantly use a self-exclusion mechanism that inactivates interference mechanisms when target RNA base pairs with the repeat region of the crRNA-guide (43). Alternatively, Type I systems use a protein-mediated mechanism to bind specific dsDNA PAM sequences (44). Here we observe that the type IV-A PAM targeting mechanism is biassed against non-complementary bases at specific positions (e.g. -2 Guanine is not complementary to the crRNA repeat, but is selected against), and prefers non-complementary bases at others (e.g. -5 Guanine is preferred), we believe these preferences support type IV-A PAM recognition mechanism that is protein-mediated and does not rely on base-pairing with the crRNA repeat.

DNA binding by a crRNA-guided Csf-complex will displace the non-target strand forming an R-loop (Figure 5A). In type I CRISPR systems, the Cas3 helicase-nuclease is recruited to the R-loop formed by the Cascade complex. Once loaded onto the non-target DNA strand Cas3 uses metal-dependent ATP binding and hydrolysis to unwind and translocate in a 3’-5’ direction from the target site, while the nuclease domain degrades displaced ssDNA (45–47). The data presented here suggest that in type IV-A systems CasDinG may play a similar role. Like Cas3, CasDinG is an ATP and metal-dependent DNA helicase. Although CasDinG unwinds with the opposite polarity as Cas3 (5’-3’ instead of 3’-5’), the helicase activity of CasDinG would still allow CasDinG to travel from the site of Csf complex targeting, perhaps modifying or destroying the DNA duplex to provide bacterial immunity with a yet to be discovered mechanism (Figure 5B). Type IV-A reliance on CasDinG helicase activity is consistent with recent work by Guo et al demonstrating that gene silencing by the *Pseudomonas oleovorans* type IV-A system appears to deplete RNA transcripts away from the site of Csf complex targeting. Interestingly, this transcript depletion appears to be bidirection suggesting that a second CasDinG can be loaded onto the target trand, perhaps through unwinding beyond the Csf complex (Figure 5C-D). Additionally, mutation of the Walker B motif impaired *in vitro* helicase activity and *in vivo* immunity, suggesting the immune system relies on helicase activity for proper function. While it is possible that ATP binding and hydrolysis could regulate another CasDinG function, we find the lines of evidence compelling enough to hypothesize that CasDinG ATP dependent DNA helicase activity is essential to type IV-A immunity.

The accessory vFeS and arch domains are essential to type IV-A immunity (Figure 4E). However, the role of these domains in type IV-A immunity remains unclear. Biochemical work with XPD and Rad3 helicases from the same family as CasDinG demonstrated that mutation of all four cysteines coordinating the FeS cluster within the FeS cluster domain disrupts helicase activity (48, 49). Additionally, mutating conserved residues on the surface of the human XPD arch domain disrupted helicase activity (38). Notably, these mutations did not disrupt ssDNA-dependent ATPase activities, indicating both the FeS and arch domains play important roles in unwinding but are not essential for ATP binding or hydrolysis. These previous studies suggest the CasDinG vFeS and arch domain deletions likely disrupted helicase, but not ATPase activities, further supporting the hypothesis that the helicase activities of CasDinG are essential to type IV-A function. However, the CasDinG accessory domains may also play additional essential roles in type IV-A immunity such as mediating interactions between the Csf crRNA-guided complex, nucleic acid substrates, and perhaps other proteins that modify DNA. We expect that a more clear understanding of the role of CasDinG in type IV-A immunity will be revealed when structures and biochemistry of CasDinG in association with the Csf complex and dsDNA targets become available.

## DATA AVAILABILITY

The NGS data from the PAM depletion assay were deposited to NCBI GEO under the accession number GSE211446. The model coordinates and structure factors for the CasDinG structure have been deposited in the Protein Data Bank under PDB code 8E2W.

## SUPPLEMENTARY DATA

Supplementary Figures 1-9

Supplementary Table 1

## ACKNOWLEDGEMENTS

We thank Randy Read and Tom Terwilliger for helpful discussions on using AlphaFold models for molecular replacement, the Chase Beisel lab for providing the PAM library and helpful discussions on performing the PAM library screen, and members of the Jackson Lab for discussions about experimental design and the manuscript.

