## Supplementary Data File for "CasDinG is an ATP-dependent 5’-3’ DNA helicase with accessory domains essential for type IV CRISPR immunity"

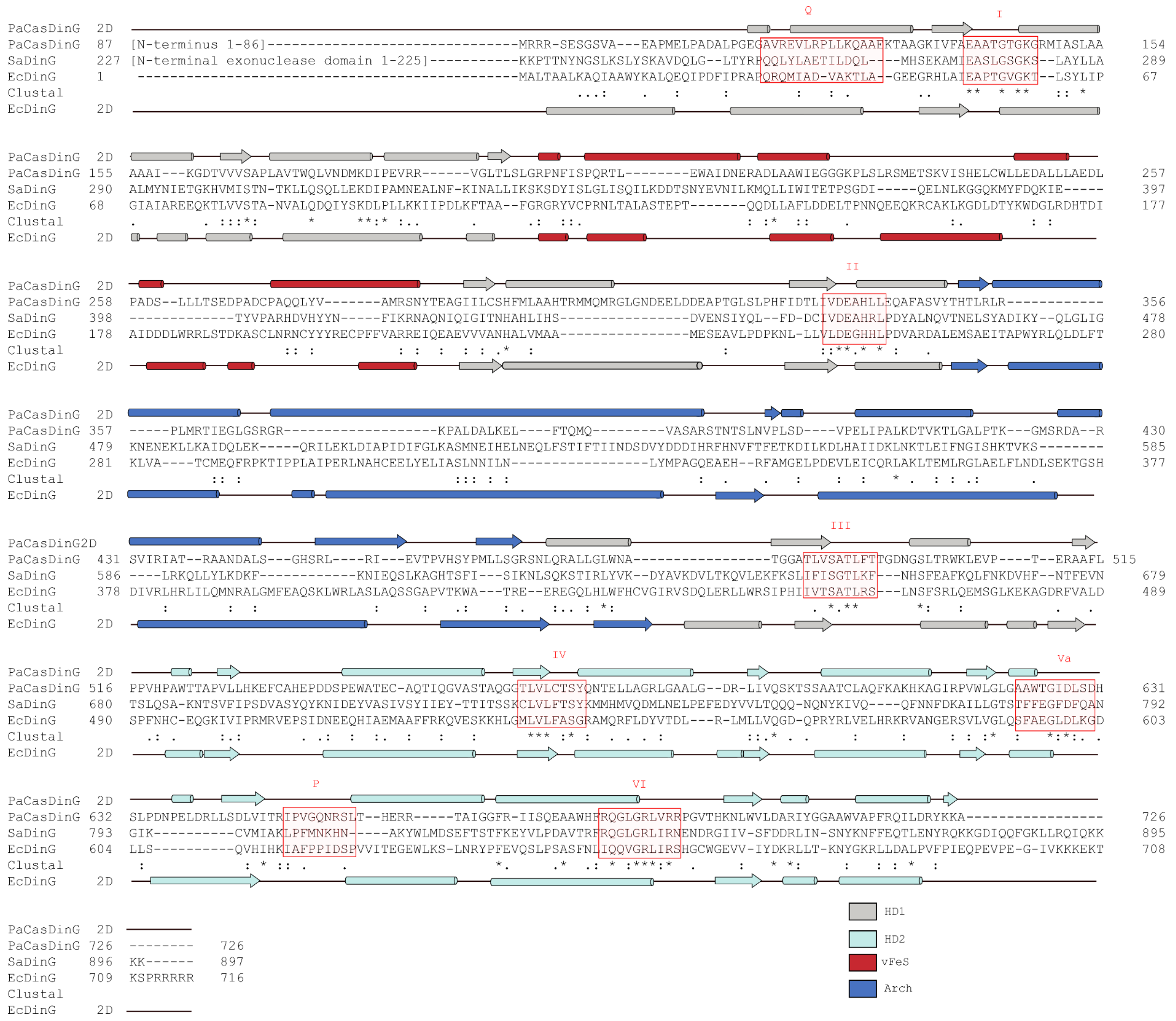


**Supplementary Figure 1. Multiple sequence alignment of *Pseudomonas aeruginosa* CasDinG, *Staphylococcus aureus* DinG, and *Escherichia coli* DinG using Clustal Omega.** Secondary structures of both Pa83 CasDinG and EcDinG are plotted above and below the sequence alignments, respectively. Conserved helicase motifs are highlighted in red.

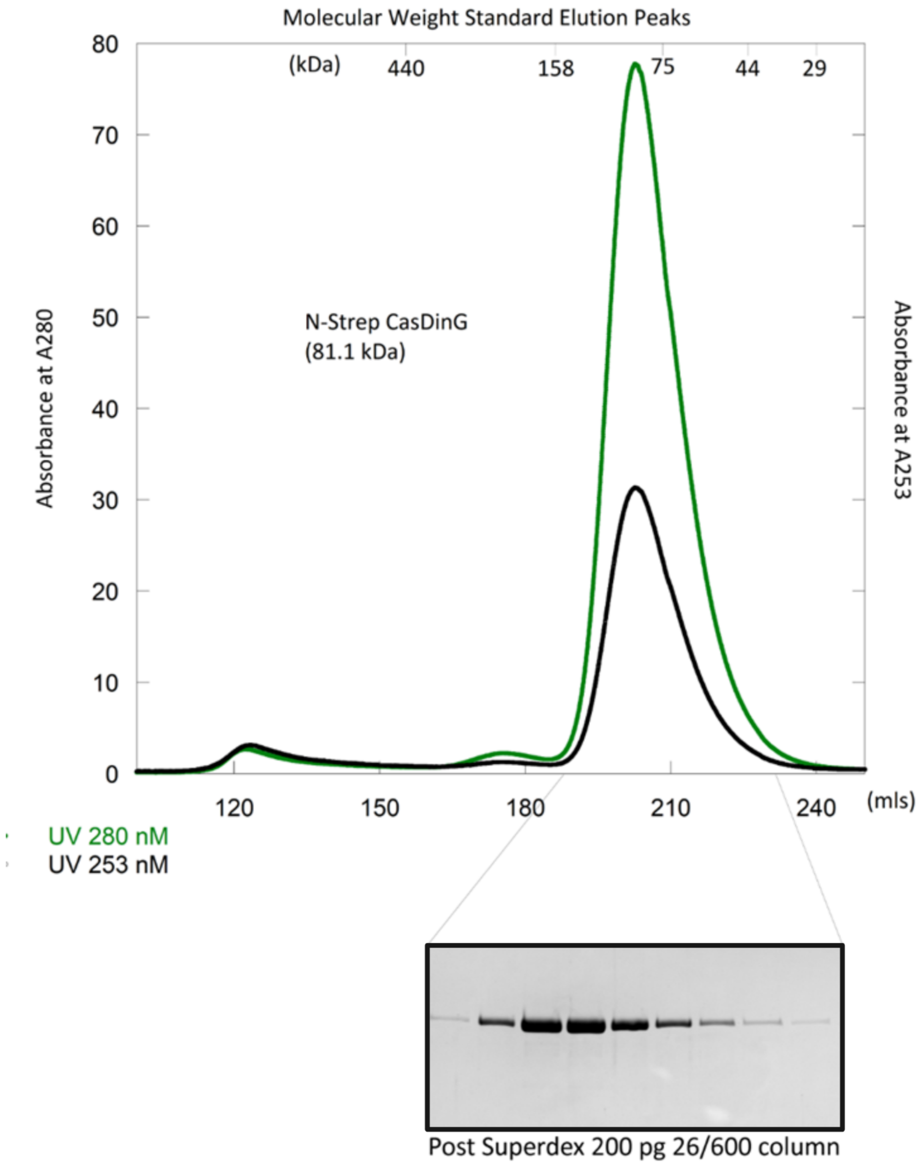


**Supplementary Figure 2. Purification of CasDinG.** Size exclusion chromatogram of N-term Strep tagged CasDinG over a Superdex 200 pg 26/600 column with the associated SDS-PAGE gel depicting protein purity.

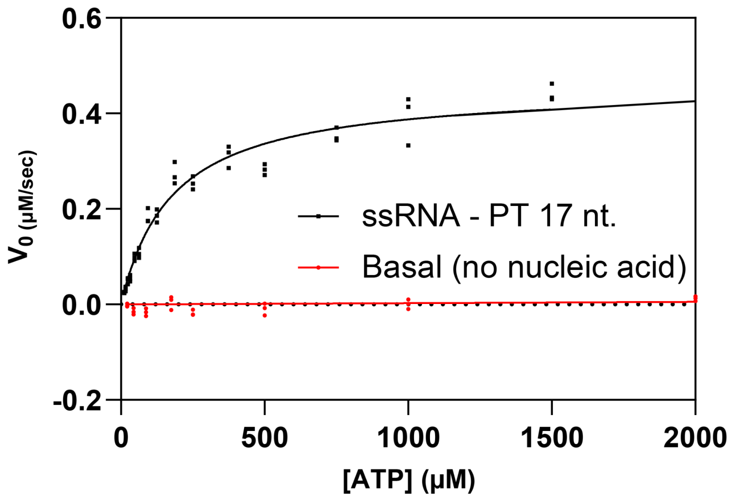


**Supplementary Figure 3. Michaelis-menten curves showing CasDinG rates of ATP hydrolysis in the presence of PT-ssRNA 17 nt. and no nucleic acid (basal).** PT-ssRNA 17 nt. data were generated using 20 nM CasDinG and 200 nM nucleic acid, resulting in a calculated kcat of 23 molecules of ATP hydrolyzed per second. Basal data was collected with 500 nM CasDinG and the resulting data could not be fit to the Michaelis-Menten equation.

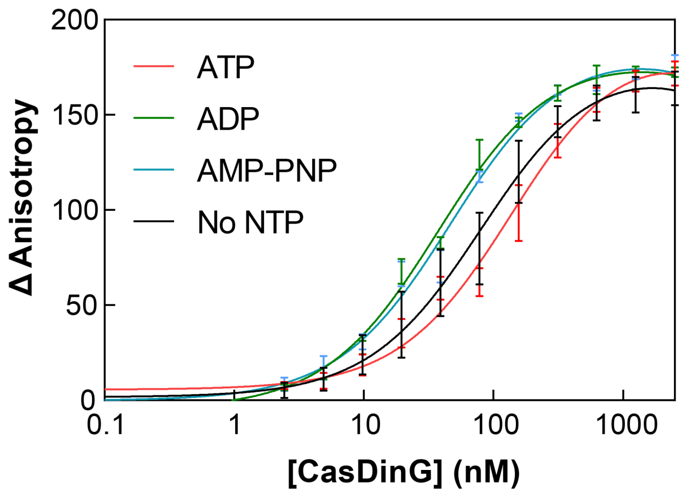


**Supplementary Figure 4. CasDinG binding to 5'-FAM labeled single-stranded DNA (17 nt.) in and out of the presence of ATP and analogs.** Binding affinities do not appear to differ substantially in the presence of ATP or nucleotide analogs.

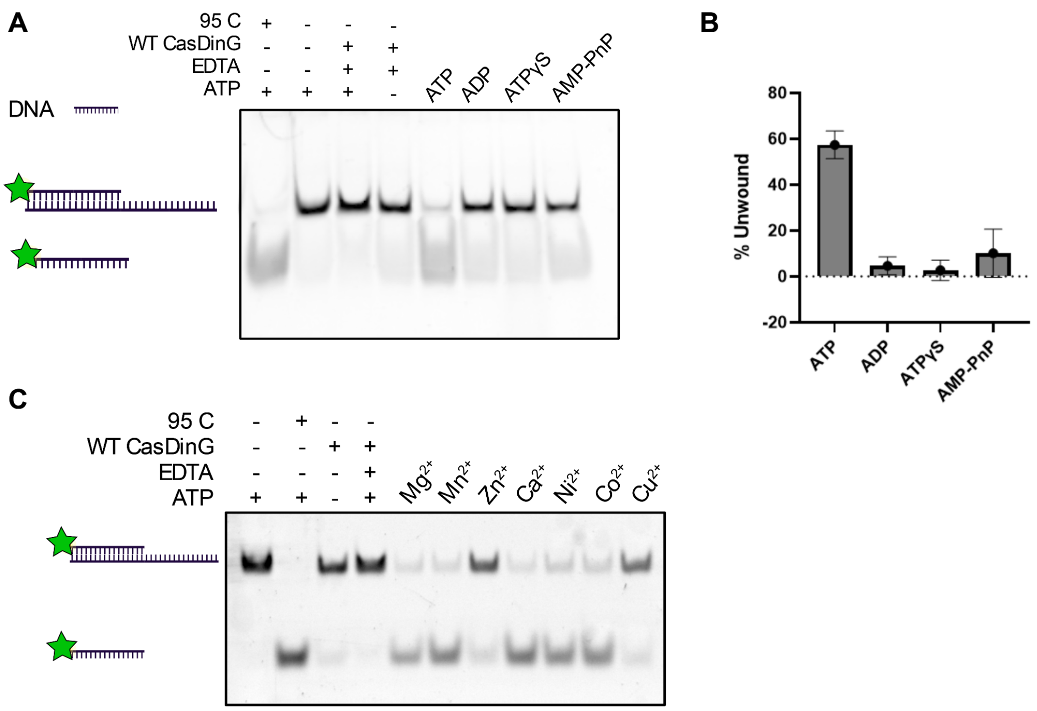


**Supplementary Figure 5. Helicase assays using ATP analogs and divalent metals.** (**A**) Unwinding of 5' FAM labeled DNA overhang duplex and ATP analogs, native PAGE shows substantial unwinding only with ATP. (**B**) % Unwound graph showing in-gel densitometry analysis of helicase assays using ATP and analogs with a 5' FAM labeled DNA overhang duplex substrate. Only helicase reactions using ATP had higher than 20% duplex unwound. (**C**) Helicase assay using 5' FAM labeled DNA overhang duplex and various divalent metals. Native PAGE shows unwinding with $Mg^{2+}$, $Mn^{2+}$, $Ca^{2+}$, $Ni^{2+}$, and $Co^{2+}$.

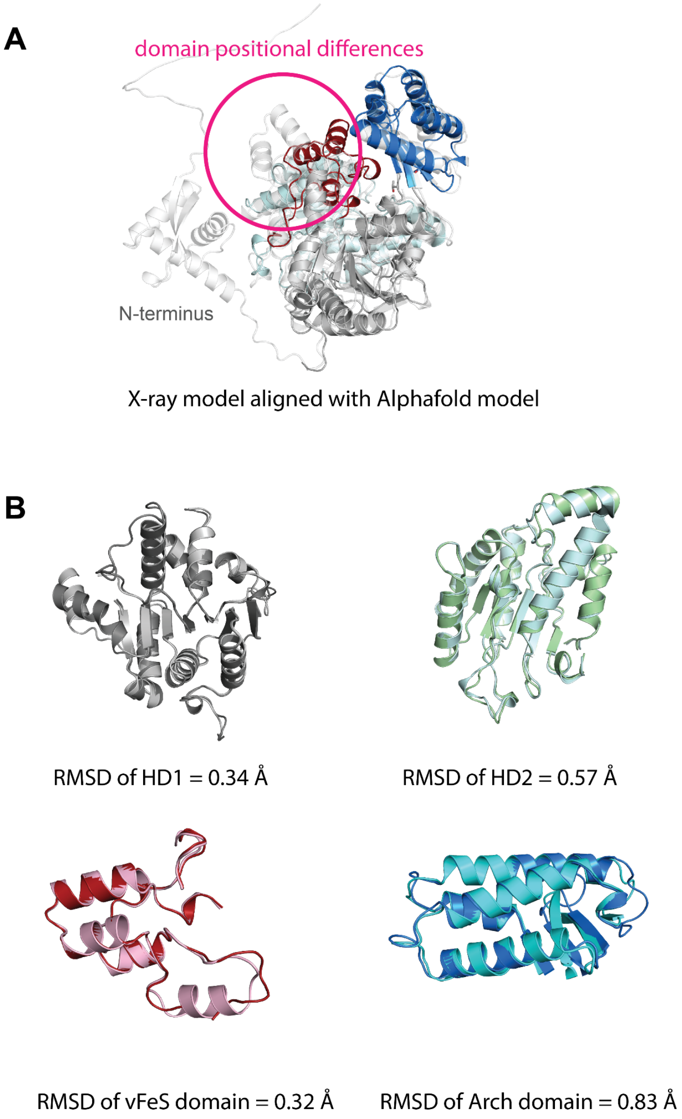


**Supplementary Figure 6. Structural alignment of X-ray model and Alphafold structure prediction model. (A**) Alignment of the x-ray crystal model and the alphafold model reveal differences in vFeS domain position. (**B**) Alignments of the HD1, HD2, vFeS and Arch domains of the x-ray model and alphafold prediction model, all alignments show RMSD of less than 1 Å.

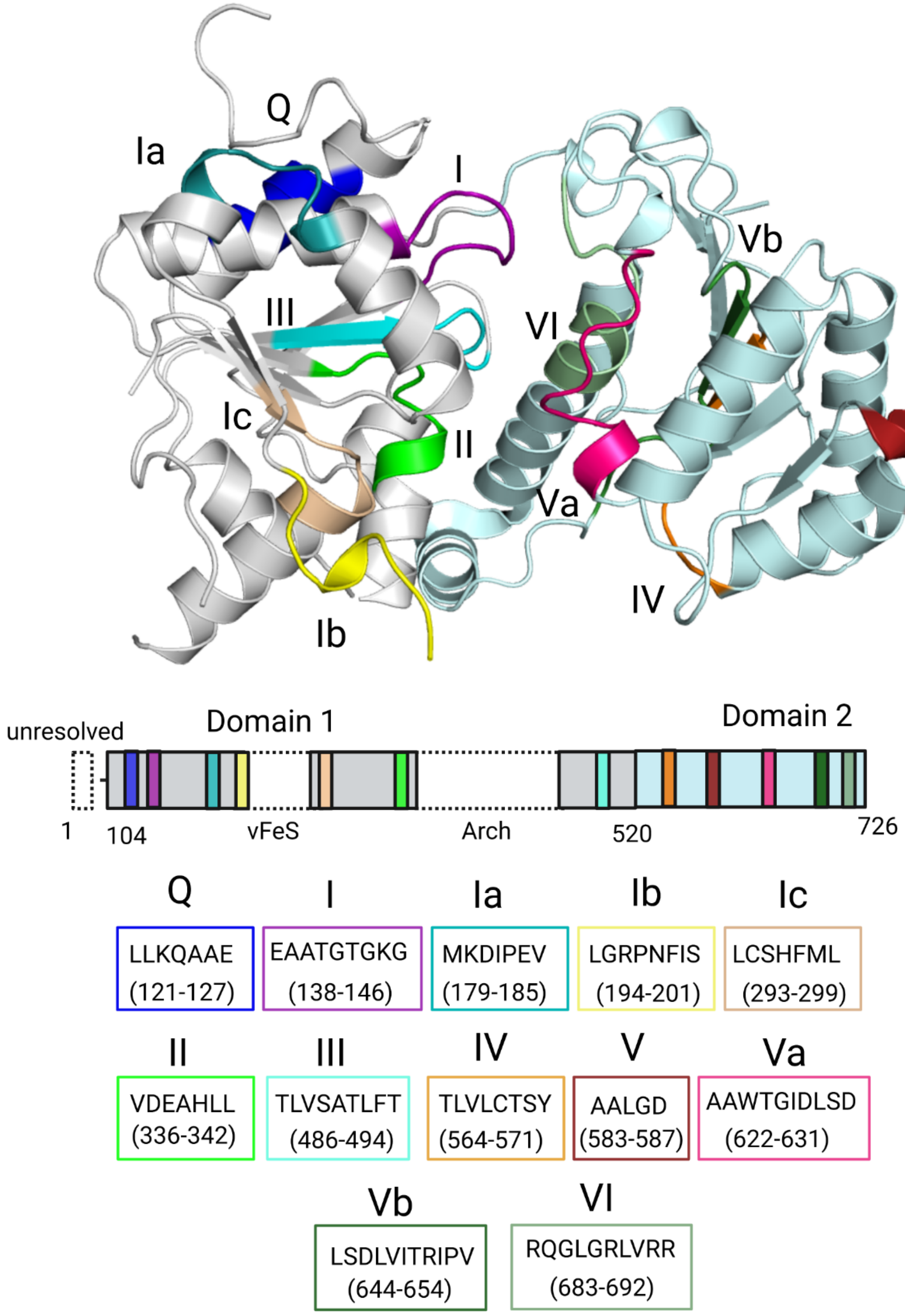


**Supplementary Figure 7. Helicase motifs of CasDinG.** Motifs Q, I, Ia, Ib, Ic, II & III are located within the helicase domain 1 whereas motifs IV, V, Va, Vb & VI are located in helicase domain 2.

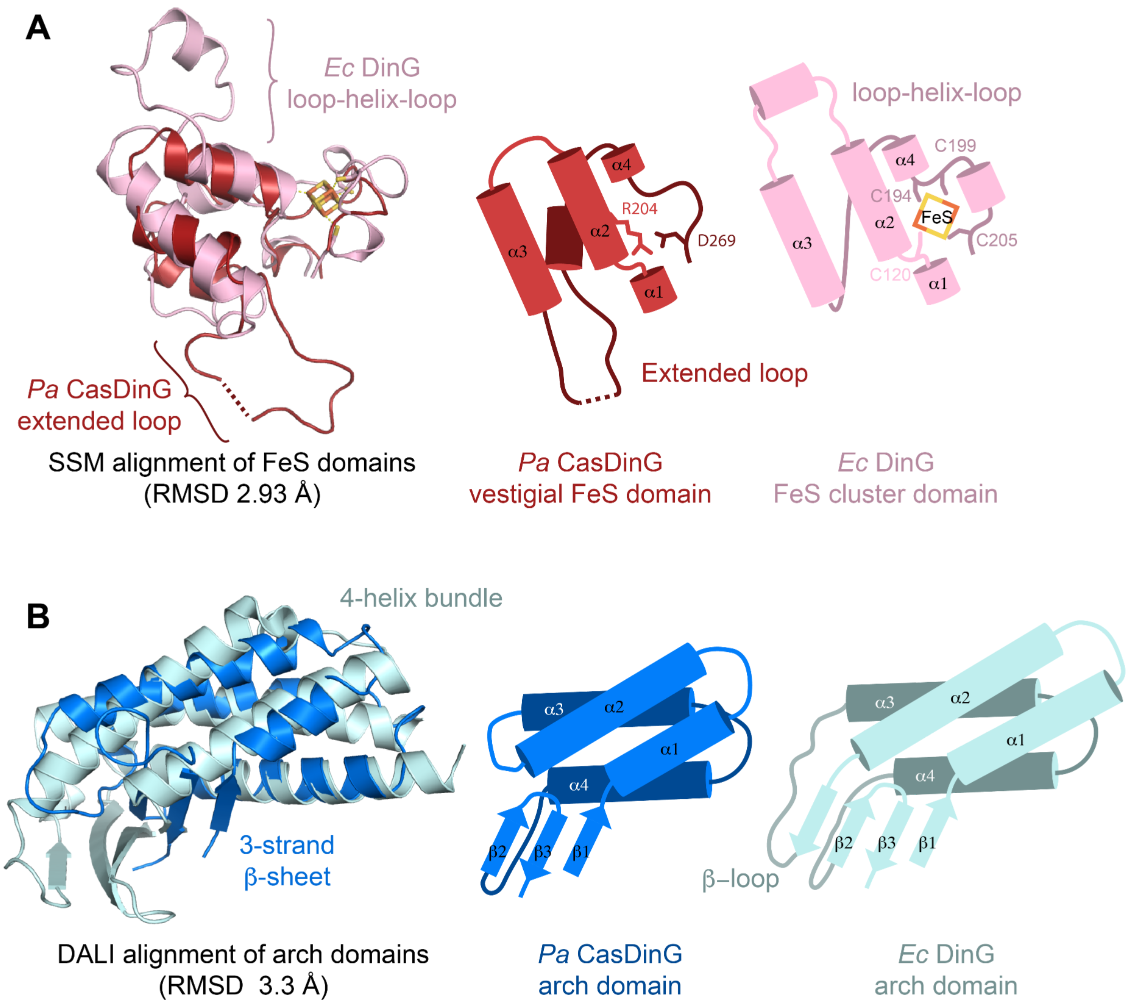


**Supplementary Figure 8. Domain alignments between CasDinG and *E. coli* DinG.** (**A**) SSM alignment of the FeS domains highlight the lack of Fe atoms in the CasDinG structure as well as the presence of an extended loop between α3 and α4. (**B**) DALI alignment of the arch domains highlights the lack of a β-loop connecting α2 and α3 in the CasDinG structure.

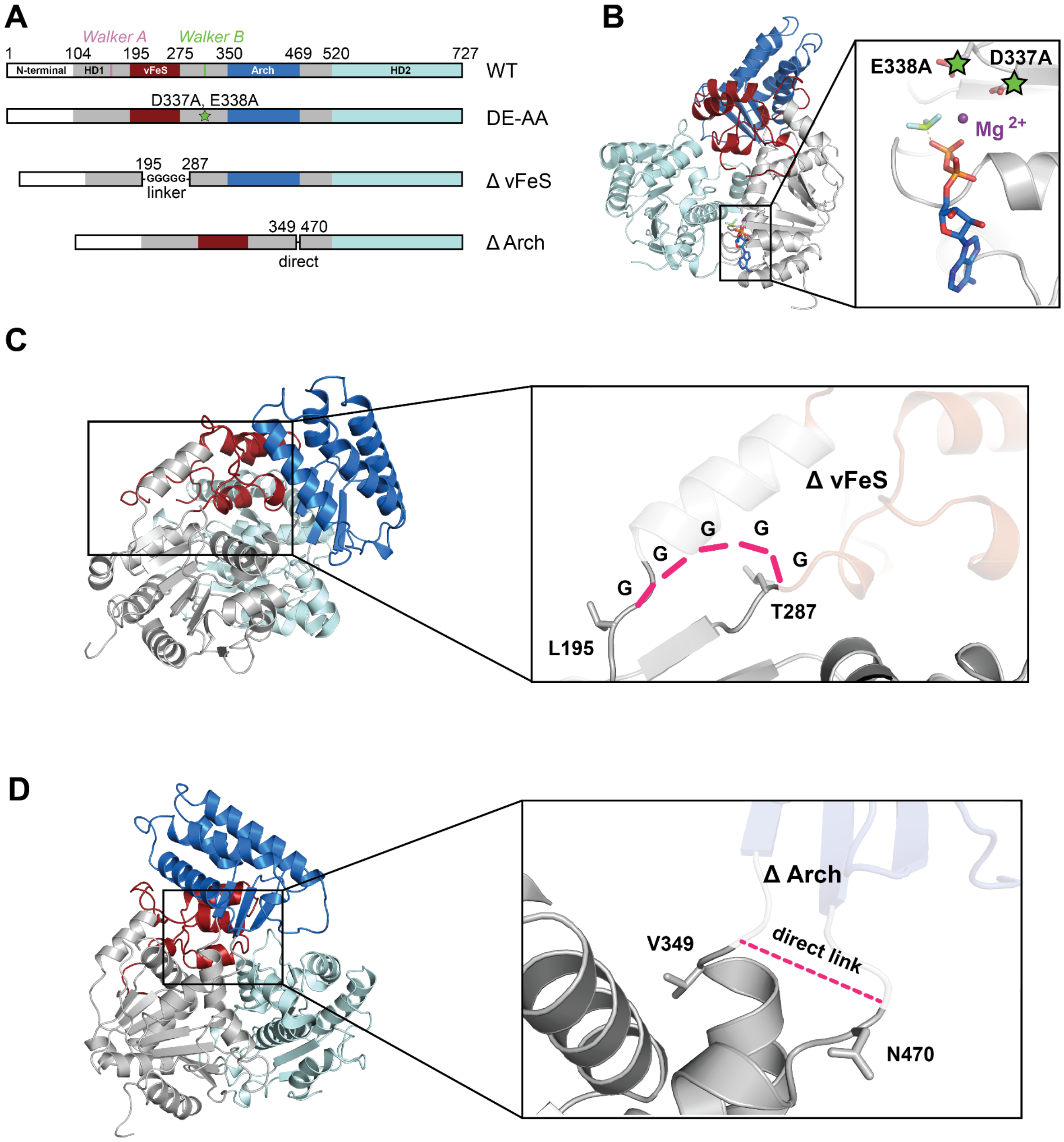


**Supplementary Figure 9. Domain mutant design.** (**A**) Schematics of the protein sequence for WT (top), the Walker B DE-AA mutant, the vFeS domain deletion, and the Arch domain deletion. (**B**) Alignment of CasDinG with ADP-BeF bound to *E. coli* DinG with the residues mutated in the Walker B mutant highlighted with green stars. (**C**) Design of the vFeS domain deletion mutant. The pink dotted line indicates how L195 and T287 are connected in the mutant. (**D**) Design of the Arch deletion mutant where V349 and N470 are directly linked.

| Oligo Identity | DNA/ RNA | Sequence | Length (nt). | Complementary Partner | Purpose |
|---|---|---|---|---|---|
| CasDinG_14 | DNA | 5’-TCGTCACCAGTACAAAC-3’ | 17 | 20,21,17,37 | Helicase Assay, ATPase, Anisotropy |
| CasDinG_17 | DNA | 5’-GTTTGTACTGGTGACGA-3’ | 17 | 14 | Helicase Assay, ATPase |
| CasDinG_20 | DNA | 5’-TTTTTTTTTTTTTTTTGTTTGTACTGGTGACGA-3’ | 33 | 14 | Helicase Assay, ATPase |
| CasDinG_21 | DNA | 5’-GTTTGTACTGGTGACGATTTTTTTTTTTTTTTT-3’ | 33 | 14 | Helicase Assay, ATPase |
| CasDinG_36 | RNA | 5’-UCGUCACCAGUACAAAC-3’ | 17 | 20, 37 | Helicase Assay |
| CasDinG_37 | RNA | 5’-UUUUUUUUUUUUUUUUGUUUGUACUGGUGACGA-3’ | 33 | 36,14 | Helicase Assay |
| CasDinG_42 | RNA | 5’-U*C*G*U*C*A*C*C*A*G*U*A*C*A*A*A*C-3’ | 17 | 43 | Anisotropy, ATPase |
| CasDinG_B2 | DNA | 5’-TCGTCACCAGTACAAACTACAACGCCTGTAGCATTCCACA-3’ | 40 | | Helicase , ATPase, Anisotropy |

**Supplementary Table 1. Oligonucleotides used in helicase, ATPase, and anisotropy assays.** *is indicative of a phosphorothioated backbone.
